# Dynamic morphoskeletons in development

**DOI:** 10.1101/657585

**Authors:** Mattia Serra, Sebastian Streichan, L. Mahadevan

## Abstract

Morphogenetic flows in developmental biology are characterized by the coordinated motion of thousands of cells that organize into tissues, naturally raising the question of how this collective organization arises. Using only the Lagrangian kinematics of tissue deformation, which naturally integrates local and global mechanisms along cell paths, we can identifying the Dynamic Morphoskeletons (DM) behind morphogenesis, i.e., the evolving centerpieces of multi-cellular trajectory patterns. The DM is model and parameter-free, frame-invariant, robust to measurement errors, and can be computed from unfiltered cell velocity data. It reveals the spatial attractors and repellers of the embryo, objects that cannot be identified by simple trajectory inspection or Eulerian methods that are local and typically frame-dependent. Computing the DM underlying primitive streak formation in chicken embryo and early gastrulation in the whole fly embryo, we find that the DM captures the early footprint of known morphogenetic features, and reveals new ones, providing a geometric framework to analyze tissue organization.

---

During embryonic development, cells undergo large scale coordinated motion during the process of tissue and organ formation that together shape the embryo. Understanding these processes requires integrating molecular, cellular and multi-cellular perspectives across a range of length and time scales, linking cellular-level gene expressions and regulatory signaling networks [1–4] to long-range intercellular interactions and mechanical force generation [5–8]. These approaches are complemented by advances in live imaging techniques [9] that allow for the detailed tracking of cellular trajectories [10–14], providing exquisite geometric and kinematic information on tissue morphogenesis. Some natural questions that these experimental approaches raise include: Can one correlate cell position, cell velocity and cell-cell interactions with cell and tissue fate decisions? Can one link gene expression levels and cellular trajectories with active force generation to help unravel the biophysical basis for morphogenesis? Can one quantitatively analyze cell motion data to predict the ultimate outcomes of tissue morphogenesis and organ development in normal and pathological situations? Here we address the last question by providing a mathematically grounded framework to determine the evolving centerpieces of morphogenetic movements using experimentally determined cellular trajectories, thus providing an important step in bridging the gap between bottom-up mechanistic approaches and top-down statistical and computational approaches [15, 16] (Fig. 1a).

**FIG. 1:**
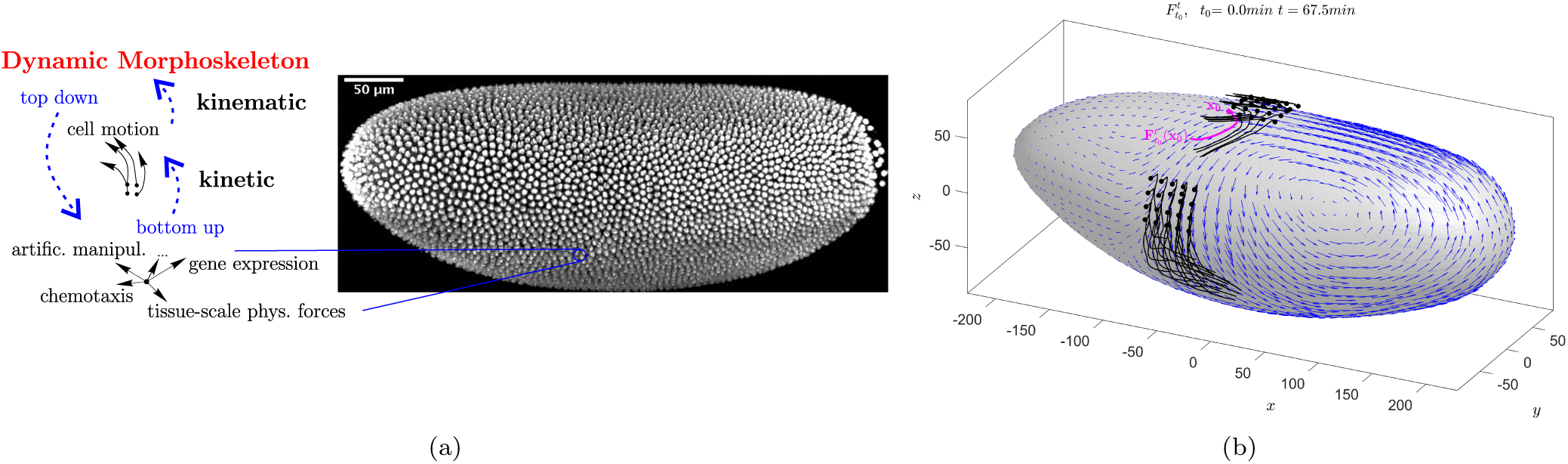
(a) Right: Drosophila melanogaster embryo during gastrulation obtained by light sheet microscopy. Left: Schematic of bottom-up and top-down approaches to study cell motion. Bottom-up approaches focus on studying local mechanisms driving cells. Top-down approaches focus on studying patterns of cell motion caused by local and global driving mechanisms. The DM uncovers the centerpieces of cell trajectory patterns in space and time. (b) Snapshot of cell trajectories (solid lines) over a time interval [*t*_0_, *t*] starting at black dots, along with the instantaneous velocity field (blue). The full trajectory evolution is available in the SI - Movie S1s.

Minimally, any framework that aims to analyze spatio-temporal trajectories in morphogenesis requires a self-consistent description of cell motion that is independent of the choice of reference frame or parametrization. This frame-invariant description of cell patterns is termed objective [17], and ensures that the material response of a deforming continuum, e.g. biological tissue, is independent of the observer. To quantify this notion, we start by considering two coordinate systems used to describe a system: the first corresponding to **x ∈** ℝ^3^, as shown in Fig. 1b, and a second one 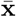 defined as

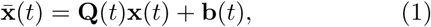

where **Q**(*t*), **b**(*t*) are a time dependent rotation matrix and translation vector. A quantity is objective (frame invariant) if the corresponding descriptions in the **x** and 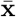 transform according to specific rules [17]. In particular, scalar quantities must remain the same 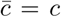, vectors must transform according to (1), and second order tensors as **Ā**= **QAQ**^⊤^. Taking the time derivative 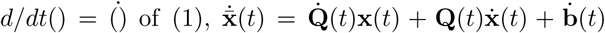, one can easily see that the velocity field and the streamlines, which are trajectories of the frozen velocity field, are frame dependent, i.e. any metrics based on them for comparative purposes are likely to be erroneous owing to the inability to remove the dependence on measurement artifacts associated with variations in the choice of reference frames etc. By contrast, the rate of strain tensor - the symmetric part of the velocity gradient - is objective [17].

Driven by the recent revolution in imaging morphogenetic flows and cellular movements [18, 19], a range of approaches have been developed to characterize mesoscopic cellular behavior. These include statistical tools based on the connectivity between neighboring sites [20], and methods quantifying cell shape changes and cell intercalation by mapping the temporal evolution of strain rates between neighboring cells [5, 6]. However, because of the general time dependence of cell motion, any velocity or velocity gradient features such as streamlines or strain rates differ substantially from Lagrangian trajectory patterns that integrate over the history of particles motion. As a concrete example, we consider the time evolution of cell trajectories (black) and instantaneous velocities (blue) of a Drosophila embryo dataset superposed on a single instantaneous final snap-shot shown in Fig. 1b (see also SI-Movie S1). If one averages the rate of strain at a fixed (Eulerian) location on the embryo **x**_0_ (magenta dot) over a time interval [*t*_0_, *t*], this average ignores completely the (Lagrangian) trajectory path (magenta line) of a cell starting from **x**_0_ at time *t*_0_. Therefore, Eulerian methods are inherently suboptimal for studying cellular flows, and suggest that a frame-invariant Lagrangian method is more suitable to assess global morphogenetic flows.

Here, we use the notion Lagrangian Coherent Structures [21], initially derived to study nonlinear dynamical systems such as those associated with complex fluid flow patterns to create an objective kinematic framework for analyzing cell motion. This allows us to uncover the *Dynamic Morphoskeleton* underlying morphogenesis-i.e., the attracting and repelling organizers of cell trajectories in space and time. We illustrate our results by studying primitive streak formation during early morphogenesis in the chick embryo and gastrulation in the fly embryo, using imaging datasets obtained by light-sheet microscopy (LSM).

## DEFINING THE DYNAMIC MORPHOSKELETON USING LAGRANGIAN COHERENT STRUCTURES

In general, trajectories of time dependent dynamical systems have complicated shapes, are sensitive to changes in their initial conditions, and are characterized by multiple spatial and temporal scales. However, underlying these complicated paths, one often finds a robust skeleton that organizes the spatiotemporal structures in the dynamical system - referred to as Lagrangian Coherent Structures (LCSs) [21], which shapes trajectory patterns and provides a simplified description of the overall dynamics. They involve information obtained by integrating the trajectories in space-time and thus serve as a memory trace of the dynamical system. They can be defined for large or small time spans [22]. In a general setting, we schematize this in Figure 2a and illustrate the impact of attracting and repelling LCSs (red curves) on trajectory patterns (black curves) over a finite time interval [*t*_0_, *t*]. The combined effect of attracting and repelling LCSs is shown in Fig. 2c. For example, blue trajectories represent two cells that were initially very close (blue dots) but end up far apart. Even though they end up far apart, and hence are apparently subject to very different fates, they end up on the same attracting LCS after separating from a repelling LCS. Therefore, assessing the system through individual trajectories, despite being Lagrangian, will return poor results.

**FIG. 2:**
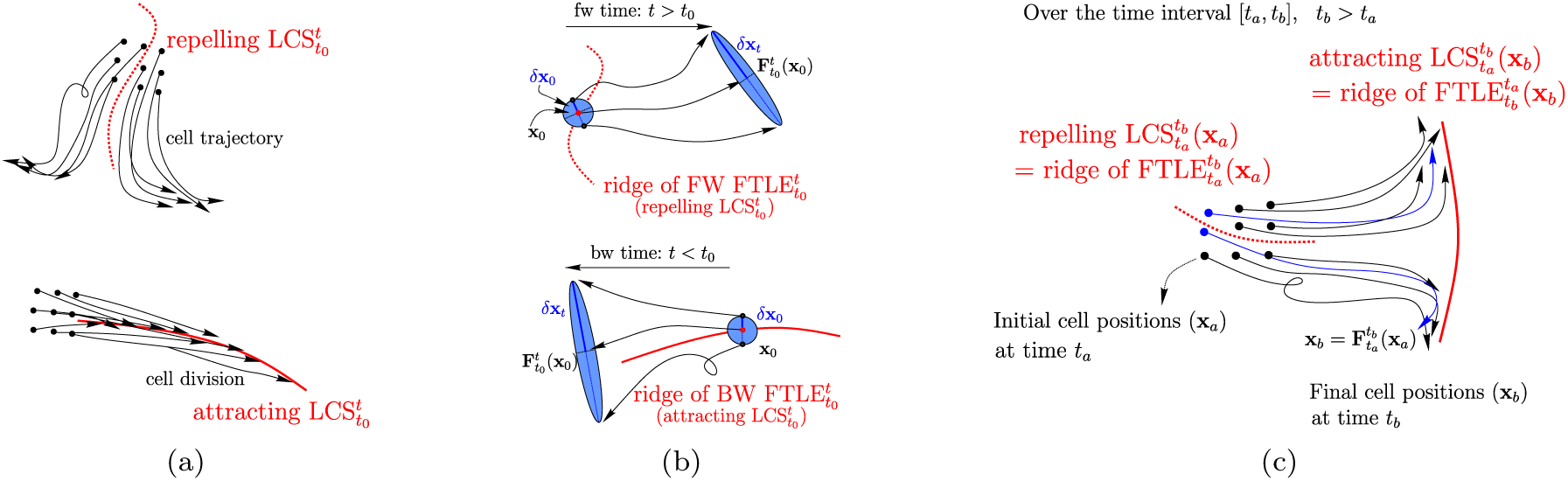
(a) Sketch of attracting and repelling LCSs (red) acting as centerpieces of cell trajectory (black) patterns over the time interval [*t*_0_, *t*]. Initial and final positions of cells are marked by circles and triangles. LCSs evolve in time and space providing the DM of cell motion. (b) The 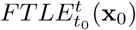 measures the maximum separation (∼|*δ***x**_*t*_*/δ***x**_0_|) induced by the flow at **x**_0_ over the time interval [*t*_0_, *t*] between two initially close points in the neighborhood of **x**_0_. A forward time FTLE ridge - a set of points with high FTLE values - marks a repelling LCS whose nearby points from opposite sided of the ridge will experience the maximum separation over [*t*_0_, *t*], *t* > *t*_0_. Similarly, a backward time FTLE ridge demarcates an attracting LCS, i.e., a distinguished curve at *t*_0_ which has attracted initially distant particles over [*t, t*_0_], *t < t*_0_. (c) Illustration of attracting and repelling LCSs over a time interval of interest [*t*_*a*_, *t*_*b*_], *t*_*b*_ *> t*_*a*_, during which cells move from their initial configuration **x**_*a*_ to their final one 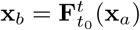. The forward FTLE is a scalar field over **x**_*a*_ while the backward FTLE over **x**_*b*_. Blue trajectories show cells that start close to each other from opposite sides of a repelling LCS, and end up far apart along the same attracting LCS. Although single trajectories assessment would assign different fates to these two cells, LCSs correctly identify their similar fate.

While there are a number of methods to determine Lagrangian (i.e., with memory) Coherent Structures [23], the Finite Time Lyapunov Exponent (FTLE), despite its limitations [21], remains the most used because it is computationally simple. The FTLE is characterized by a scalar field used to locate regions of high separation (or convergence) of initially close (distant) particles over the time interval [*t*_0_, *t*]. Denoting by **v**(**x**, *t*) a time dependent velocity field obtained from imaging data, the induced Lagrangian flow map 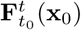 is given by

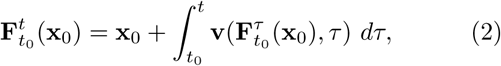

which maps the initial positions (of cells, membranes or nuclei, for example) **x**_0_ at time *t*_0_ to their final positions at time *t.* The FTLE is then defined as

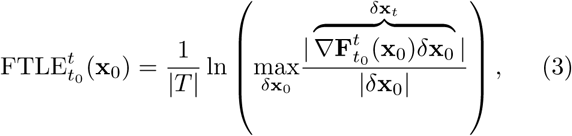

where |·| represents the absolute value and ∇ the Jacobian with respect to **x**_0_. The FTLE is thus a measure of the maximum separation rate between a trajectory starting at **x**_0_ and a neighboring one starting at **x**_0_ + *δ***x**_0_ (in the limit of small |*δ***x**_0_|), over the time interval [*t*_0_, *t*] (Fig. 2b) (see SI for explicit formulas for computing (3)).

We note that the FTLE depends on the base time *t*_0_, the spatial location **x**_0_ - which correspond to the positions of Lagrangian particles at the base time - and the final time *t*, which sets the time scale *T* = *t* − *t*_0_. As illustrated in Fig. 2b, a set of points **x**_0_ with high forward FTLE values (FW FTLE ridge) marks a region at *t*_0_ whose neighboring particles from opposite sides of the ridge will get repelled achieving maximum separation at the later time *t* = *t*_0_ + *T, T >* 0. Similarly, a backward FTLE ridge marks regions that at the base time *t*_0_ have attracted initially distant particles over the time interval [*t*_0_ + *T, t*_0_], *T <* 0 (Fig. 2b). Together, the forward and backward FTLE fields associated with varying time scales *T* uncover the exact spatial locations of repelling and attracting LCSs, along with the times at which they appear and cease to exist. We further note that over a time interval of interest [*t*_*a*_, *t*_*b*_], *t*_*b*_ *> t*_*a*_ during which cells move from their initial configuration **x**_*a*_ to their final one 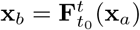, the forward FTLE is a scalar field over **x**_*a*_ while the backward FTLE is a scalar field over **x**_*b*_. Therefore, over the time interval [*t*_*a*_, *t*_*b*_], trajectories initially at opposite sides of FW FTLE ridges will be repelled from each others and get attracted to BW FTLE ridges by time *t*_*b*_ (Fig. 2c).

A mechanical interpretation of (3) follows by noting that the 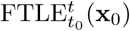 is proportional to the logarithm of the highest eigenvalue of the Cauchy–Green strain tensor 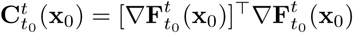, a naturally invariant measure of deformation of a continuous medium. Hence it represents the maximum deformation induced by the flow over [*t*_0_, *t*] on an infinitesimal area element centered at **x**_0_ (Fig. 2b), and thus provides an exact link between the DM and the Lagrangian strain experienced by cells during morphogenesis. Separation or convergence of cell trajectories captured by the FTLE can arise from a combination of isotropic (volume or area) changes – due e.g., to cell divisions and ingression – and anisotropic (shear) deformations – due to cell shape changes. In Fig. 2b, the former effect is shown by the changes in the light blue areas between circles and ellipses, while the latter is shown via the circle-to-ellipse deformation which can occur while keeping the area unchanged. To distinguish between these two effects, we compute the isotropic volume shrinkage induced by 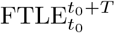 as

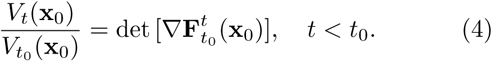

This scalar field is based at the final configuration **x**_0_ and integrates information over the time interval [*t, t*_0_]. It attains values greater than unity in regions where initially far trajectories converge because of isotropic contraction, and is less than unity in regions where initially close trajectories diverge due to isotropic expansion. Since volume/area changes and shape changes completely characterize all possible deformations in a continuous medium, equations (3-4) completely characterize tissue deformations during morphogenesis. We now deploy these concepts on two paradigmatic problems in large scale morphogenetic flows: primitive streak formation in the chick embryo and gastrulation in the whole fly embryo. In both cases, we will follow the spatiotemporal evolution of the DM in terms of the FTLE fields as a function of their memory *T*, and thus determine the attracting and repelling manifolds underlying tissue organization.

## RESULTS

### Primitive streak formation in chicken embryo

The primitive streak (PS) is a hallmark of bilateral symmetry in many organisms and involves large scale cell flows to form an axial structure that serves to organize embryogenesis. The formation of this structure is best understood in the chick embryo and involves coordinated flow of more than 100,000 cells in the epiblast. Here we use a cell velocity dataset of a Myr-GFP embryo obtained by a dedicated LMS as described in [24]. The velocity field is given on a uniform rectangular grid of size [5.17*mm* ×1.66*mm*] over a time interval of approximately 13*h* from the freshly laid egg (stage EGXIII), prior to the onset of tissue movement, with spatial resolutions of 0.65*µm* and temporal resolution of 3*min*. As in [24], we filtered the cell velocities using a centered averaging filter with a 5 × 5 spacial, and a ∑ 5 time instances temporal window sizes. We then compute attracting and repelling LCSs as backward and forward FTLE (Supplementary Material, Methods) for a set of time scales |*T*| spaced by 15*min*, and spanning the full length of the dataset.

The top panel of Fig. 3a shows the backward 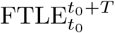 for the complete dataset: the large values highlight the attracting LCS demarcating the formed PS towards which cells (green dots) got attracted after |*T*| = 13*h* from the freshly laid egg (0*h*). The bottom panel shows the forward 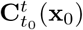 for the same |*T*| : the large values highlight the position of a strong repelling LCS at the freshly laid egg time from which close cells will separate over the next 13*h*. While embryonic regions towards which cells tend to cluster have been studied before [16, 24], our analysis also shows the presence of undocumented repelling regions that are complementary to the attractors. Remarkably, cells starting at the opposite sides of the repelling LCS, oriented perpendicularly to the AP direction, move towards opposite sides even within the PS (see SI-Movie S2). Green dots indicate the position of cells at time *T*, and started at time 0 from a uniform rectangular grid. The FTLE values have units *min*^*-*1^ and the axis are in *µm*. In the white regions, the FTLE is unavailable because cells starting there have left the domain in which velocity is available within the time interval [*t*_0_, *t*_0_ + *T*]. (See SI-Movie S2) shows the full evolution of FW and BW FTLE fields for different *T*, along with cell positions, uncovering the DM behind PS formation. The PS is characterized by a first converging phase towards a vertical structure perpendicular to AP and located at *x* ≈3600*µm*, followed by an AP extension leading to the final PS shown in the top panel of Fig. 3a. Remarkably, already within 60*min*, while cells barely moved, the BW FTLE already shows a footprint of the PS forming perpendicularly to the AP direction encircled by a blue ellipse in Fig. 3c. The random looking distribution of the FTLE on the left side of the figure is typical of either random or uniform (deformation-less) motions. We note that differently from existing studies, where the early location of the PS is obtained by following backward in time the cells belonging to the formed PS [24], our approach does not use future data, hence revealing the footprint of PS formation only from the Lagrangian tissue straining field.

**FIG. 3:**
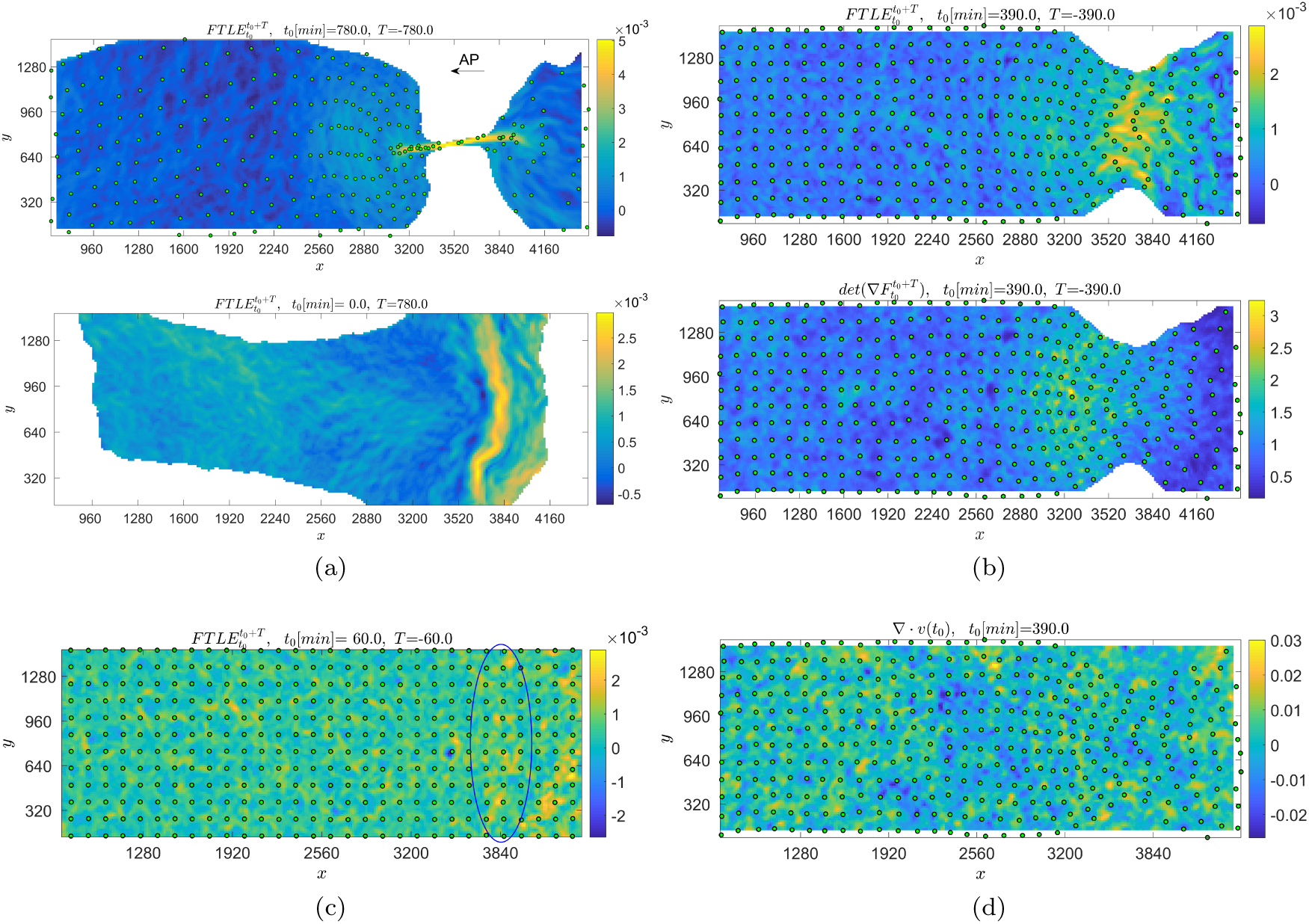
(a) Top: BW FTLE corresponding to the full dataset (|*T*| = 13*h* from the freshly laid egg stage) highlights the attracting LCS corresponding to the (PS). Cell positions at 13*h*, initialized from a uniform gird at 0*h*, are shown by green dots. White areas corresponds to regions where the FTLE is unavailable because trajectories left the domain over which the velocity field is defined. Bottom: FW FTLE corresponding to the full dataset highlights repelling LCSs. The FTLE has unit min^*-*^ 1, the axis units are in *µm* and posterior is at larger *x* values. The full time evolution of BW and FW FTLE for different *T*, along with the corresponding cell positions, is available as SI-Movie S2. (c) BW FTLE ridge (attracting LCS) for *T* = 1*h* highlights the early footprint of the PS (blue ellipse) using only data within [0, 1]*h* during which cells (green dots), initially released on a uniform rectangular grid, barely moved. The visually random distribution of FTLE on the left side of the figure is typical of either random or uniform (deformation-less) motions. (b) Top: BW FTLE ridge (attracting LCS) for |*T*| = 6.5*h*. Bottom: Isotropic area shrinkage (dimensionless) as described in (4) for the same time interval of the top panel. Higher values (*>* 1) denote embryo regions at 6.5*h* which isotropically shrank over [0, 6.5]*h*. The PS captured by the BW FTLE ridge shows that anisotropic deformations dominate isotropic ones within the first 6.5*h*, shifting the mild isotropic attraction ridge towards the posterior pole. (d) The divergence (in min^*-*1^) of the instantaneous velocity field at time 6.5*h*. Negative values demarcate regions of isotropic instantaneous contraction rate. This Eulerian field – which is the instantaneous limit of the Isotropic area shrinkage shown in panel (b) Bottom – misses both the current location of the forming PS, and the clustering regions due to only Lagrangian isotropic contraction. Green dots show cell positions at 6.5*h*, started at 0*h* from a uniform rectangular grid.

As noted above, clustering of cells arises from the combination of anisotropic deformation and isotropic volume shrinkage along cell trajectories. The top panel of Fig. 3b shows the BW FTLE, as in Fig. 3a, for a shorter time scale |*T*| = 6.5*h*, and the bottom panel shows the clustering contribution due to isotropic area shrinkage over the same time interval, as described by (4). Comparing the two panels we observe that anisotropic deformation plays a determinant role in the PS formation over 6.5*h*, as its combined effect with isotropic shrinkage shifts the attracting LCS towards the posterior side compared to the clustering mild ridge revealed by looking only at isotropic deformation. In Fig. 3d we show the instantaneous divergence of the velocity field at |*T*| = 6.5*h* after freshly laid eggs as in in Fig. 3b, along with current cell positions. The divergence field is the instantaneous version (*T* →0) of the finite-time area shrinkage described by (4), and negative values demarcate regions characterized by instantaneous isotropic area contraction rate in *min*^*-*1^. Remarkably, the instantaneous divergence field shows no signature of the actual location of the forming PS (Fig. 3b Top), as well as of the finite-time isotropic area contraction (Fig. 3b Bottom), highlighting the need of Lagrangian methods for studying morphogenesis.

While filtering velocities is necessary for Eulerian methods, owing to their sensitivity to noise and measurement errors, it may hide finer-scale structures whose characteristic sizes are smaller than the filtering window sizes. By contrast, Lagrangian methods are intrinsically more robust because the integration of cell velocities along trajectories acts like a filter. In Supplementary Fig S3 and Supplementary Movie 1, we show the same analysis of Fig. 3a using the raw unfiltered velocity. We find that the DM is exceptionally robust to noise and measurement errors, and is perfectly computable from raw data without ad-hoc filtering of cell velocities.

### Gastrulation in the fly embryo

Instead of focusing only on a specific morphogenetic feature, here we analyze the early development of the entire fly embryo. During gastrulation of Drosophila, about 6000 cells on the embryonic blastoderm on the embryo surface undergo global morphogenetic flow which induce severe tissue deformation, finally giving rise to the three germ layers. We compute the DM on an “ensemble averaged” cell velocity dataset from 22 wild-type Drosophila melanogaster embryos undergoing gastrulation, developed in [25]. Each velocity dataset is obtained combining in toto light sheet microscopy [11, 12] and tissue cartography [26], and consists of coarse-grained velocities averaged with a spatial window of approximately 5 cell size. The velocity field is given on 1800 grid points over the fixed apical embryo surface (Fig. 1b), and covers a time interval of approximately one hour with a temporal resolution of 75*sec*, starting well before the ventral furrow formation when the earliest flows are characterized by a dorsal sink and a ventral source. Because of cell moving on a curved surface, the FTLE depends on the metric tensor describing the curviness of the apical embryo surface. In the Supplementary Material we provide the exact formula for computing FTLE as well as the Cauchy–Green strain tensor 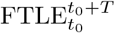 characterizing the finite time tissue deformation induced by cell motion along arbitrary curved surfaces.

We compute the DM for a set of time scales |*T*| spaced every 1.5*min*, and spanning the full length of the dataset. Fig. 4a shows the backward 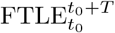 corresponding to the full extent of the dataset, whose high values highlight the position at time 67*min* towards which cells get attracted over the time interval [0, 67.5]*min*. The FTLE has unit *min*^*-*1^ and the axis units are *µm* (SI-Movie S3) shows the full evolution of a 3D view of BW FTLE field for different *T*, whose final time frame is shown in Fig. 4a or Supplementary Fig. 4a. On the ventral side, the initial source at the ventral pole give rise to te ventral furrow (VF) which terminates with the anterior midgut invagination and a Y-shaped bifurcation close to the posterior pole (top panel of Fig. 4a). The inset shows the VF of the embryo imaged using LSM. While it is not surprising that the VF corresponds to an attracting LCS, his formation and connection to the initial ventral source is inaccessible to both Eulerian methods and simple inspection of cell trajectories. At later times compared to VF formation, a U-shaped attracting LCS (mid panel of Fig. 4a) emanating in proximity of the posterior pole demarcates the lateral boundaries of the germband extension. On the dorsal side, drosophila gastrulation is characterized by several transverse structures which include the cephalic furrow (CF), the anterior and posterior folds, and the posterior midgut invagination. Given the coarse-grained nature of the velocity field, structures whose width is smaller than 5 cell size, such as the CF and the transverse folds, should be invisible to cell trajectories obtained integrating the velocity dataset. Yet, we find a sharp transverse attracting LCS (mid and bottom panel of Fig. 4a), arising from the initial dorsal sink, and continuously moving towards the A pole. The coexistence of such transverse attracting LCS, the U-shaped one, and the VF clearly explain cell trajectory patterns shown in SI-Movie S4, which is the same as SI-Movie S3 along with cell positions. Supplementary Fig. 4b shows the last snapshot of SI-Movie S4.

**FIG. 4:**
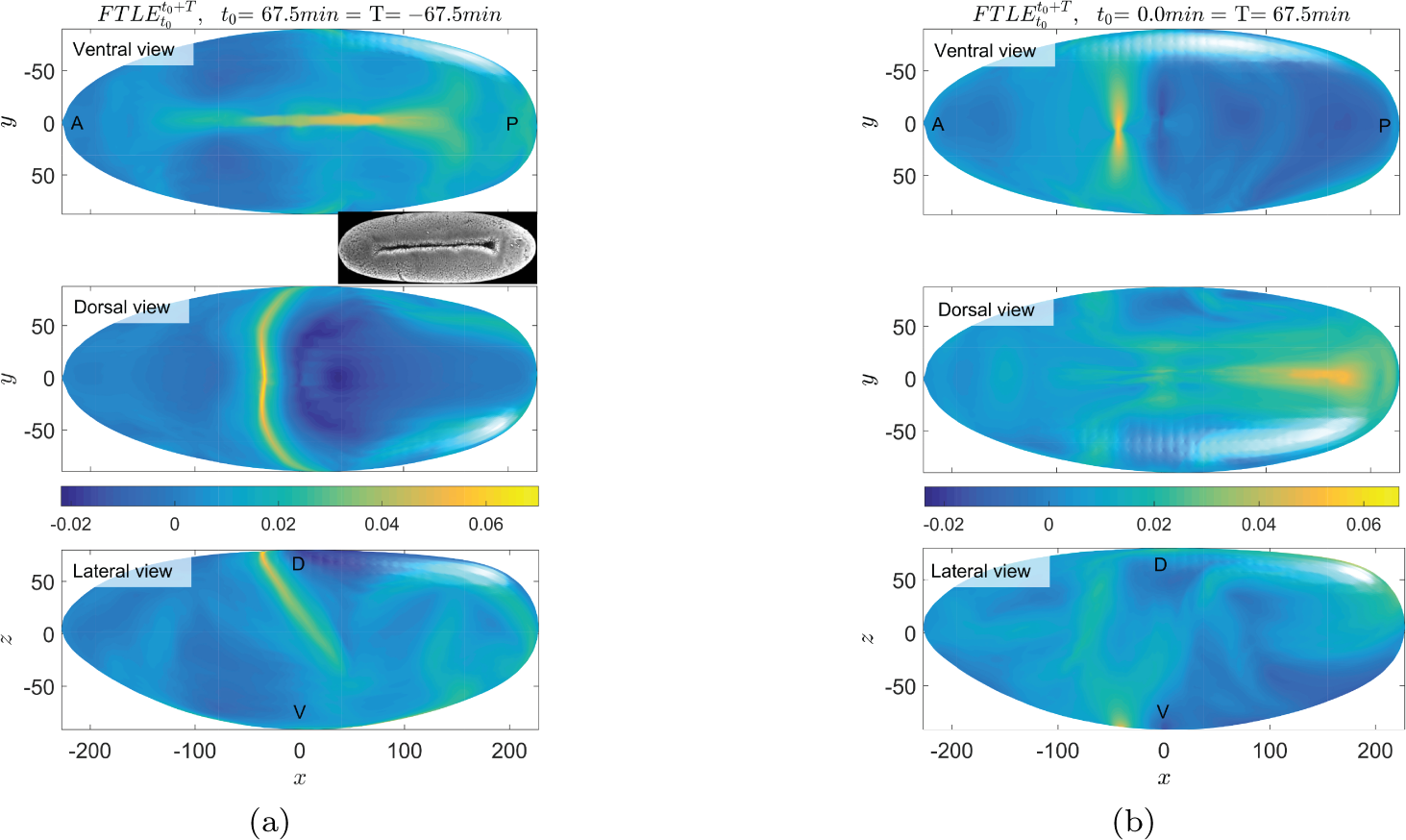
(a) BW FTLE corresponding to the full extent of the dataset (|*T*| = 67.5 min) highlights the attracting LCSs during early gastrulation of Drosophila. The top panel shows a strong attracting LCS capturing the VF, which terminates on the A side with the anterior midgut imagination and on the P side with a Y-shaped bifurcation. The inset shows the VF from an embryo image obtained by LSM. The mid and bottom panels show a strong transverse attracting LCS shifted towards the A pole and a U-shaped milder attractor emanating from the P pole region. A 3D embryo view with the full time evolution of BW FTLE for different *T* is available as SI-Movie S3, SI-Movie S4 is the same as Movie S3 along with cells position in green. (b) FW FTLE corresponding to the full extent of the dataset highlights repelling LCSs. The two main repellers are a transverse ridge on the ventral side that separates the anterior region from the rest of the embryo, and a dorsal ridge in the AP direction which appears synchronously with the VF, and repels cells towards the lateral sides. The FW FTLE evolution for different *T* is available as SI-Movie S5. The FTLE has unit min^*-*^ 1 and the axis units are in *µm*.

Figure 4b shows repelling LCSs (forward 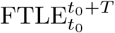) corresponding to the full extent of the dataset, whose high values highlight the position at time 0*min* from which close cells separate over the interval [0, 67.5]*min* (SI-Movie S5) shows the full evolution of a 3D view of the FW FTLE field for different *T*, whose final time frame is shown in Fig. 4b or Supplementary Fig. 4c. The DM reveals that the initial dorsal sink gives rise to a repelling LCS along the AP axis which appears synchronously with the VF and repels cells towards the lateral sides, while the initial ventral source evolves into a repelling LCS, perpendicular to the VF, which moves towards the anterior and separates the A region from the rest of the embryo. Together, SI-Movie S4 and SI-Movie S5 show the evolution of attracting and repelling LCSs in space and time fully characterizing the DM. The above results show that the DM can either coincide with known morphogenetic features as the VF, or reveal new attracting and repelling structures which invariably shape trajectory patterns as a result of both local and global mechanisms.

## CONCLUSIONS

Using only available kinematic data associated with cell trajectories, we have provided a simple but rigorous kinematic framework for analyzing morphogenetic flows to uncover the evolving centerpieces of cell trajectory patterns which we term the Dynamic Morphoskeleton (DM). The DM is frame invariant and based on a Lagrangian description of tissue deformation captured by the FTLE, which naturally combines local and global mechanisms along cell trajectories. The DM is robust to noise and measurement errors, can be computed directly from raw data without filtering of cell velocities, and thus can find fine-scale structures whose sizes are smaller than typical filtering window sizes. The DM is composed of attracting and repelling LCSs towards which cells converge to, or diverge from, over a specific time scale. Of particular note is evidence not just for attracting regions, but repelling regions that are just as important in determining cell fate (e.g., cells starting at opposite sides of a repelling LCS typically undergo different fate).

We have found that DM can either coincide with known morphogenetic structures, or reveal new ones which invariably shape trajectory patterns. The DM allow us to uncover the early footprint of morphogenetic features as well as their evolution, which remain inaccessible to Eulerian methods and simple inspection or cell trajectories. For example, in the chick PS formation, we have found that already within 1*h* from freshly laid egg, the DM identifies the cells that will be part of PS. At the same time, we have found the presence of a repeller that separates cells even within the PS.

Since the DM is driven solely by kinematic information, it can be computed directly from cell motion data and is agnostic to the mechanisms generating them. This is both an advantage and a disadvantage - as it provides a framework to study the organizers of development, and yet does not shed light on their origin. Connecting the DM to known gene expression patterns and mechanical processes is a natural next step to understand the relative importance of different co-existing spatiotemporal mechanisms. Independently, our methods might also be useful to compare different morphogenetic phenotypes, and bridge the gap between bottom-up and top-down modeling approaches to morphogenesis.

## ACKNOWLEDGMENTS

We thank Kees Weijer for helpful discussions and for providing the chicken embryo dataset.

